# Signatures of parallel evolution in sperm-mediated paternal effects in threespined sticklebacks

**DOI:** 10.64898/2026.08.26.747353

**Authors:** Jennifer K. Hellmann, Miles Bensky, Alison M. Bell

## Abstract

1. Transgenerational plasticity (TGP)- when parental environments influence offspring phenotypes - is ubiquitous across taxonomic groups and can have benefits for offspring beyond what is possible with developmental plasticity, particularly when selective pressures are high early in life. However, patterns of TGP vary widely across populations and species, and the evolutionary processes shaping this variation remain poorly understood.
2. Here, we tested whether repeated evolutionary transitions result in parallel or population-specific evolutionary divergence in TGP relative to ancestral conditions. We examined sperm-mediated paternal effects across two ancestral marine and three derived freshwater populations of threespined stickleback fish (*Gasterosteus aculeatus*). We exposed fathers to dragonfly larvae (endemic to freshwater) or sculpin (endemic to all populations) predators and measured both paternal response to predators as well as antipredator behavior and growth in larval offspring.
3. Fathers behaviorally responded to the presence of sculpin predators, but not dragonfly larvae. However, we found strong paternal effects in response to both predators in all populations. Further, the magnitude of TGP did not differ between marine and freshwater populations, suggesting that TGP does not become genetically accommodated as marine populations move into freshwater habitats.
4. We found some evidence consistent with parallelism in both within and trans-generational plasticity: 1) personal exposure of larval stickleback to dragonfly larvae elicited strong antipredator responses in freshwater populations that were absent in marine populations, and 2) paternal predation exposure consistently increased offspring growth in marine populations while slowing growth in freshwater populations. In contrast, paternal effects altered offspring behavior in population-specific ways, with strong sex-specific effects of paternal exposure emerging in response to endemic predators.
5. Adaptive evolution is a two-step process, in which heritable genotypic and phenotypic variation must first be present and then selected on. Therefore, high population-level variation in TGP suggests the capacity for rapid evolution of parental effects, while signatures of parallelism and sex-specific patterns suggest that TGP may evolve in targeted ways in response to ecological stressors.

## Introduction

Organisms frequently encounter spatial or temporal fluctuations in their environment that elicit adaptive responses via genetic adaptation and/or phenotypic plasticity. Transgenerational plasticity (TGP) occurs when the environment experienced by one generation influences the phenotypes of future generations (A. M. Bell & Hellmann, 2019). TGP is nearly ubiquitous across taxonomic groups (Yin et al., 2019), can improve offspring fitness (e.g., in response to parental exposure to predation risk (Storm & Lima, 2010), pathogens (Moore et al., 2019), toxins (Cong et al., 2019), rising temperature (Donelson et al., 2016)), and can have benefits for offspring beyond what is possible with developmental plasticity, particularly when selective pressures are high early in life (Donelan et al., 2020). Given the importance of TGP for fitness, there is increasing interest in understanding factors that shape its evolution and whether population (Heckwolf et al., 2020; Vijendravarma & Kawecki, 2015; Walsh et al., 2016) and species-level (Ramos-Muñoz et al., 2024; Sultan et al., 2009) variation in TGP reflects adaptive responses to different ecological conditions.

One way to assess whether TGP adaptively evolves is to compare patterns of TGP among populations of the same species that vary in their evolutionary histories with different ecological stressors. Theory predicts at least three non-exclusive signatures of adaptive TGP across replicate populations of the same species. First, the strength of TGP may differ between ancestral and derived populations: adaptive plasticity in ancestral ‘stem’ populations may become genetically assimilated (lost) or accommodated in derived populations (flexible stem model of adaptive evolution) (West-Eberhard, 2003). For example, gene expression due to cold stress is canalized in derived European populations of fruit flies, which facilitates adaptation to temperate environments (von Heckel et al., 2016). This may be a common outcome of adaptive radiation as selection via genetic change refines existing plasticity to produce adaptive traits that match the new environment (Levis & Pfennig, 2016). Second, if TGP is adaptive, then we expect to see signatures of parallelism in TGP: plasticity evolves in a consistent direction in response to predictable environmental variation. For example, populations of *Silene uniflora* plants show parallel changes in gene expression in response to adaption to zinc (Wood et al., 2023). This suggests TGP has the capacity to produce convergent evolutionary outcomes in the slope of reaction norms. Finally, if TGP is adaptive, TGP may be fine-tuned in response to endemic stressors: a population’s evolutionary history with a specific ecological condition influences the direction and/or adaptive nature of the plastic response. For example, parental exposure to drought increases offspring biomass in plants from dry environments, but reduces biomass in a closely related species that grows in moist environments (Sultan et al., 2009). Adaptive responses can be refined by selection in populations where stressors are endemic, while novel cues produce maladaptive or variable responses in offspring because parents have not evolved the appropriate cue-response systems (Donelan et al., 2020). We can assess evidence for these three signatures of adaptive TGP by systemically evaluating patterns of TGP across multiple replicate populations in response to both endemic and novel stressors.

Threespined stickleback (*Gasterosteus aculeatus*) are an excellent system for exploring the evolution of TGP. Sticklebacks are ancestrally marine and marine populations are relatively unchanged since the last glaciation (M. A. Bell & Foster, 1994). In contrast, numerous freshwater populations have repeated diverged from marine populations, and have become locally adapted to selective pressures in freshwater habitats (M. A. Bell & Foster, 1994), including predation risk (Wund et al., 2015). We compared the transgenerational consequences of paternal exposure to predation risk across 5 populations: two ancestral marine populations (Resurrection Bay and Rabbit Slough), two derived freshwater populations that were colonized hundreds to thousands of years ago (Cornelius and Big Beaver), and one newly established derived freshwater population (Cheney). We raised the F0 parent generation in the laboratory under common garden conditions and then exposed fathers to sculpin predators (endemic to marine and freshwater populations) or dragonfly larvae (endemic to only freshwater) immediately prior to reproduction. In stickleback, prefertilization exposure of fathers to predation risk can be transmitted to offspring via sperm/seminal fluid and alter offspring behavior and morphology (Afseth et al., 2022; Chen et al., 2021; Hellmann, Bukhari, et al., 2020; Hellmann et al., 2021; Hellmann & Rogers, 2024; Rogers & Hellmann, 2025). We observed fathers’ behavioral responses to both predators, used their sperm to generate offspring, and examined larval offspring for traits related to predation defense, including growth and antipredator behavior in response to both sculpin and dragonfly larvae. To increase power to detect TGP, we genetically sexed offspring to account for potential differences in how sons and daughters responded to paternal cues (Hellmann, Bukhari, et al., 2020; Hellmann, Carlson, et al., 2020).

Leveraging this powerful experimental design, we first assessed if ancestral and derived populations differ in their evolved responses to sculpin and dragonfly larvae (independent of paternal treatment). Population-level differences in the capacity of individuals to adjust anti-behavior behavior (within-generational plasticity) would suggest that predation risk is a selective pressure producing heritable differences in plastic reaction norms. We then tested for three signatures of adaptive evolution of TGP. First, we tested the flexible stem hypothesis by comparing TGP in marine and freshwater populations in response to a common predator (sculpin). We predicted that, if TGP becomes genetically accommodated in derived populations, this would result in divergence between marine and freshwater populations in the degree of TGP, particularly in response to sculpin predators that are common to all populations (e.g., marine populations show TGP while freshwater populations do not). Second, we tested for parallel evolution of TGP by comparing patterns of TGP among freshwater populations. We predicted that if TGP is adaptive, it would result in parallel evolution of TGP across the different freshwater populations (i.e., TGP evolves in a predictable direction as marine populations colonize novel freshwater environments). Finally, by comparing TGP in response to sculpin (endemic to all populations) and dragonfly larvae (endemic to freshwater populations), we tested for evidence that TGP is fine-tuned in response to endemic, but not novel, predators. We predicted that transgenerational responses to endemic predators (dragonfly larvae in freshwater populations) would be more adaptive than transgenerational responses to novel predators (dragonfly larvae in marine populations). Although we did not explicitly test fitness, adaptive TGP could be reflected in more rapid growth (to outgrow the gape of the predator) [26] or evidence that paternal exposure to dragonfly predators enhances offspring responsiveness to dragonfly larvae in freshwater, but not marine, populations (predator-specific paternal priming). Alternatively, it is also possible that selection on TGP could refine reaction norms, such that there is high interindividual variation in response to novel dragonfly in marine populations, while reaction norms in freshwater populations have consistent slopes (Stein & Bell, 2019).

## Methods

### Field collections

In June 2017, we collected adult threespined sticklebacks from 5 different sites in Alaska: Resurrection Bay, Rabbit Slough, Cornelius, Big Beaver, and Cheney. Cornelius and Big Beaver were naturally colonized, likely hundreds to thousands of years ago after the last glacial maximum, while Cheney was experimentally seeded in 2009 from Rabbit Slough (M. A. Bell et al., 2016). Sculpin are not endemic to Cheney, but are endemic to all other populations, including Rabbit Slough; therefore, individuals from Cheney have encountered sculpin recently in their evolutionary history. Resurrection Bay and Rabbit Slough are weakly genetically differentiated from each other (F_ST_ = 0.0076 (Hohenlohe et al., 2010)) while the freshwater populations are more strongly genetically differentiated from each other (Mike Bell, Krishna Veeramah, personal communication). We generated embryos in the field using previously established protocols (Wund et al., 2015) and shipped them to University of Illinois, Urbana155 Champaign.

### F0 parent rearing conditions

We reared all populations under common garden, freshwater conditions. Upon arrival, we incubated fertilized eggs in a cup with a mesh bottom over an air bubbler until hatching (8-13 days post fertilization). We reared fry in 9.5L (32 x 21 x 19 cm) tanks, with each clutch housed in a separate tank. We kept all families on two recirculating flow-through racks, with particulate, biological, and UV filters. We pseudo-randomly assigned family tank position such that all populations were evenly distributed across racks and shelves. We maintained fry on a 12 L : 12 D photoperiod at 15.5° ± 1°C and fed them newly hatched brine shrimp for 2 months before transitioning to a mix of frozen bloodworms (*Chironomus* spp.), brine shrimp (Artemia spp.) Mysis shrimp, and Cyclop-eeze, fed *ad libitum* daily.

Once fish were reproductively mature (June 2018), we moved fish from their individual clutch tanks into larger stock tanks and individually marked them with two elastomer marks on their dorsal side to indicate family of origin. In July– October, we switched adults to a summer photoperiod (16 L : 8 D) and placed males in individual 26.5L tanks (36L x 33W x 24H cm), visually isolated from the other males’ tanks with opaque partitions. Each tank contained two plastic plants, a sandbox, and algae to encourage nest building, and had a 12-sqaure grid (4 wide by 3 high) on the front of the tank to be used for behavioral observations.

Once they built nests, we exposed males either to a model sculpin for 1 minute every other day (21cm in length) or a model dragonfly larvae (5.1cm in length) for 1 minute every other day. Consistent with previous studies, we used a short stressor to avoid potential habituation to predation risk (Dellinger et al., 2018). Further, by beginning the treatment when males were transferred to a nesting arena, we sought to mimic the change in predator regime that males may encounter as they move into a different habitat to nest. The number of predator exposures depended on the availability of gravid females, and ranged from 4-14 times (mean 6.2 ± 2.2 s.d.), and was matched between dragonfly and sculpin-exposed fathers as much as possible. We left the males in the control group in the tank for the same amount of time as the predator exposed fish and also tapped gently on the surface of water for the same amount of time. For each male, we conducted behavioral observations on up to three of these exposures (n=94 males observed, n=268 observations total). All observations were conducted on the first through fifth exposure, with the majority (n=229) conducted on the first three exposures. For each observation, we measured activity (number of total squares visited) and time in the bottom third of the tank for three minutes before and after the simulated predator attack (or control tapping).

The day after the last exposure, we removed the male, extracted his testes, and used his sperm to fertilize the eggs of a gravid female from the same population (females originated from a different clutch to avoid inbreeding). As much as possible, we used a split clutch design: one female’s eggs were split three ways, such that each female’s eggs were fertilized by a control male, a dragonfly-exposed male, and sculpin-exposed male. The fertilized eggs were placed in a cup with mesh bottom above an air bubbler in a 9.5L tank and monitored for mold before hatching. All clutches were fed newly hatched brine shrimp upon hatching and offspring were assayed between 3-4 weeks after hatching. We assayed offspring from a total of n=95 fathers: n=18 fathers from Resurrection Bay, n=17 fathers from Rabbit Slough, n=24 fathers from Big Beaver, n=18 fathers from Cornelius, and n=18 fathers from Cheney. To control for genetic variation, we had brothers from the same family sire offspring across each of the three treatment as much as possible. Across the 5 populations, fathers were pulled from a total of n =45 starting clutches that were generated in the wild.

### Offspring behavior assay

We conducted open field assays to test the effects of paternal predation risk on offspring behavior. We used a round opaque container (radius = 7 cm) that was half-way filled with tank water and divided into eight sections circularly, with a small pebble in each section. To minimize the stress response from being caught, the fish was caught with a clear cup and gently transferred to the container, where it habituated for 15 minutes. Once 15 minutes had passed, we measured the total number of sections visited as a proxy for activity and the amount of time the fish spent within a body length to the wall while it poked perpendicularly toward the wall (thigmotaxis) using a stopwatch for 3 minutes. Thigmotaxis, or behaviors in which individuals remain close to the wall and search for escape, are often interpreted as a measure of anxiety behavior (Maximino et al., 2010). We then simulated a model predator attack with either a miniature clay sculpin model (8.1 cm in length) or a model dragonfly larvae (5.1cm in length) for approximately 5 seconds. We used both predator types to understand whether paternal exposure to a given predator primes offspring to be more responsive to that predator (e.g., offspring of sculpin-exposed fathers respond more strongly to sculpin while offspring of dragonfly-exposed fathers respond more strongly to dragonfly); assays were split relatively evenly between the two predator types (n=467 assays with a dragonfly predator, n=483 assays with a sculpin predator). This simulated predator attack often elicited freezing behavior, which is an antipredator defense. We measured the amount of time the fish spent frozen; after the fish resumed its movement, we again measured the total number of sections visited and the duration of thigmotaxic activity for 3 minutes. We scored assays live, using four different observers. Upon completion of the assay, larvae were sacrificed in an overdose of MS-222, measured for standard length (tip of the nose to the base of the caudal fin) and mass, and preserved in ethanol in a −20 freezer. We used these tissues samples to sex the fish using a genetic marker (Peichel et al., 2004). We assayed a total of 950 offspring across the 5 populations (up to 20 fish per clutch, with 10 fish per assay predator; see Supplementary Table 1 for more details on sample sizes).

### Statistical analysis

#### Paternal response to predator exposure

We used a principal components analysis (R package factoextra (Kassambara & Mundt, 2017)) to combine activity and time spent on the bottom of the tank (Spearman rank correlation: ρ=-0.54, p<0.001). Behaviors were scaled and centered, and we included two datapoints per individual for behavior before and after the simulated predator attack. We extracted one principal component with an eigenvalue of 1.50 that captured 74.8% of the variation, with larger values indicating individuals who were more active and spent less time on the bottom of the tank (‘bold’). We used MCMC generalized linear mixed models (R package MCMCglmm (Hadfield, 2010)) with a Gaussian distribution and a weak prior on the variance (V=1, nu=0.002). We included fixed effects of paternal treatment (control, dragonfly, sculpin), population, observation period (before or after the attack), day of exposure (e.g. first versus third exposure), and experimental date, to control for the fact that populations varied slightly in the time at which they reproduced within the experiment. We also included random effects of clutch ID of the father and individual father ID, to account for repeated observations. We tested for interactions of time period with all other fixed effects, as well as interactions between population and paternal treatment; we removed all non-significant interactions. We removed 2 outliers to normalize the residuals.

#### Offspring phenotypes

We ran separate models for each population to determine significant effects of paternal treatment in each population. We did this for two reasons. First, while we found no evidence for treatment by population interactions for our paternal data (Supplementary Table 2), we found statistically significant two-way interactions between paternal treatment and population for all offspring traits examined (Supplementary Table 3). This suggests that these transgenerational effects are highly variable across different populations of sticklebacks. Second, for some models, we had specific predictions about three-way interactions within a population (see below); running one model for all populations together would have created the possibility of an uninterpretable 4-way interaction.

Because we found evidence of heteroskedasticity of the residuals within some populations for some traits, we used MCMC generalized linear mixed models (R package MCMCglmm (Hadfield, 2010)) with a weak prior on the variance (V=1, nu=0.002) to analyze behavioral (activity, thigmotaxis, freezing behavior) and morphological (standard length, body condition) traits. We ran models for 200,000 iterations, with a burn-in of 3000 iterations, thin = 3, with a Poisson distribution for behavioral traits and Gaussian distribution for morphological traits.

For all behavioral models, we included fixed effects of paternal treatment (control, dragonfly-exposed, sculpin-exposed), offspring sex (male or female), offspring standard length, and assay predator (dragonfly or sculpin); for the activity and thigmotaxis models, we included two additional fixed effects of observation period (before or after the simulated predator attack) and an interaction between assay predator and observation period to understand if there was a greater change in behavior before and after the simulated predator attack depending on whether offspring encountered a sculpin or dragonfly larvae during the assay. For all behavioral models, we included random effects of observer identity, maternal family, and paternal family; for activity and thigmotaxis, we also included additional random effects of fish ID to account for repeated observations on the same individual. For the body condition model, we regressed mass on standard length and used the residuals as the dependent variable. For the length and body condition models, we included fixed effects of paternal treatment and offspring sex, with random effects of maternal family and paternal family. For length, we included an additional fixed effect of age; for body condition, we included an additional fixed effect of date, to control for seasonal effects on growth. We removed one high outlier from the standard length dataset and 2 high outliers from the mass dataset.

We tested for several additional interactions and removed interactions when they were not statistically significant. For all models, we tested for interactions between paternal treatment and offspring sex to understand whether sons and daughters responded differently to paternal treatment. In the behavioral models, we tested for a two-way interaction between assay predator and paternal treatment (freezing) and a three-way interaction among assay predator, paternal treatment, and time period (activity, thigmotaxis) to understand whether paternal exposure to a given predator may have primed offspring to respond more strongly to that predator during the assay (e.g., offspring of sculpin-exposed fathers freeze longer in response to sculpin compared to dragonfly larvae, while offspring of dragonfly-exposed fathers freeze longer in response to dragonfly). Finally, in activity and thigmotaxis models, we tested for an interaction between time period and paternal treatment, to understand if paternal treatment caused offspring to show a greater or reduced change to a simulated predator attack.

### Animal welfare note

All methods were approved by Institutional Animal Care and Use Committee of the University of Illinois at Urbana-Champaign (protocol ID 18080), including the use of a model predator. We attempted to minimize stress to experimental animals by using short-lived stressors, handling them gently (e.g., moving fish in a cup without removing them from water), and by providing refuges in the home and testing tanks.

#### Replication statement

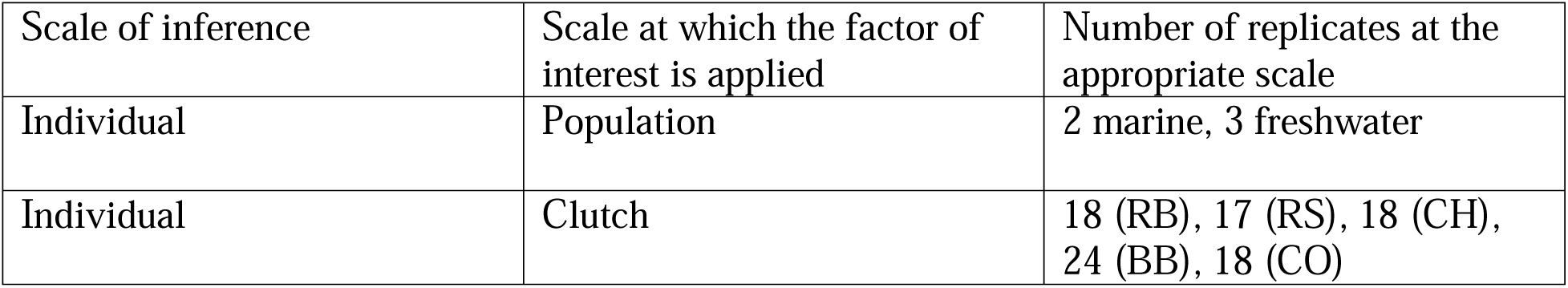

## Results

### Both fathers and offspring show strong, conserved behavioral responses to sculpin and dragonfly predators

We first assessed if ancestral and derived populations differ in their evolved responses to sculpin and dragonfly larvae. Fathers (n = 268 observations across n=94 males) showed a greater reduction in boldness (reduced their activity and increased time on the bottom of the tank) after exposure to sculpin compared to the control (time period by sculpin predator: 95% CI [-0.79, −0.12], p=0.008), while behavioral changes in response to dragonfly did not differ from the control (time period by dragonfly predator: [-0.61, 0.07], p=0.10; Figure 1). This suggests that fathers perceive the sculpin to be a greater threat than the dragonfly larvae, perhaps because sculpin prey on both adults and juveniles while dragonfly prey only on juvenile stickleback.

**Figure 1:**
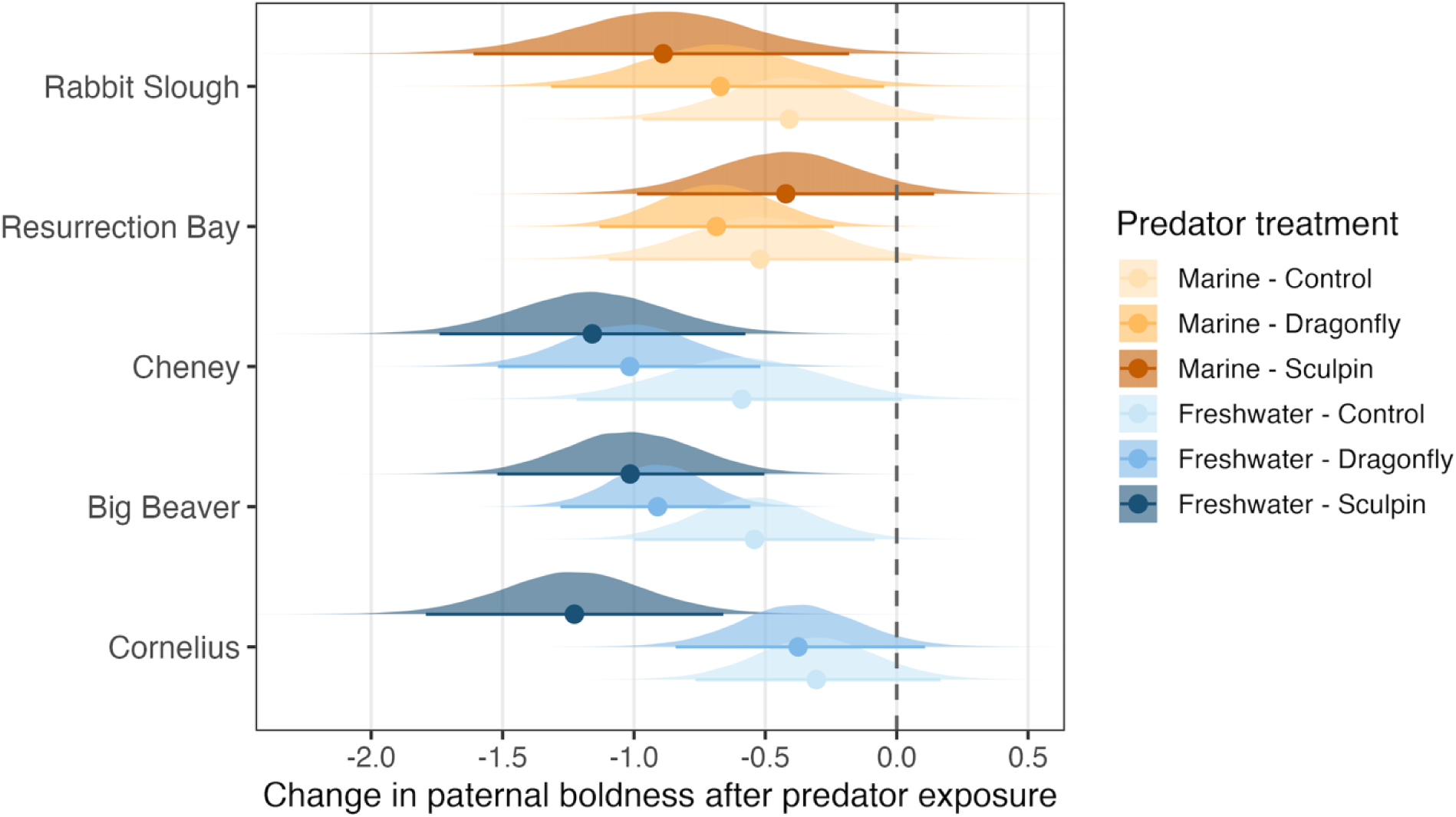
We observed paternal behavior before and after a simulated predator attack by the sculpin (dark) or dragonfly (middle) predator or a sham control (splashing on top of the tank; light). Although fathers were less bold after all exposures (0 represents no change in behavior, with negative numbers indicating less activity after the simulated predator attack), fathers showed a greater decrease in activity in response to sculpin predators relative to the control condition. There was a similar pattern with dragonfly larvae, although there was no significant difference in behavior between control and dragonfly treatments. Data shown are the posterior distributions of the MCMCglmm models, with the median (dot) and the 95% Highest Density Interval (bar).

Larval offspring generally showed significantly less thigmotaxis behavior and were less active after the simulated predator attack compared to before (significant effect of time period; Table 1), suggesting that they behaviorally responded to the simulated predator attack in the assay. However, larvae from all three freshwater populations responded more strongly to the dragonfly larvae compared to the sculpin (assay predator by time period interaction; Table 1; Figure 2) while larvae from both marine populations did not differ in their response to sculpin versus dragonfly larvae (Table 1; Figure 2). Further, fish from one marine population (Resurrection Bay) and one freshwater (Big Beaver) population spent more time frozen in the assay after encountering dragonfly larvae compared to sculpin (Table 1). Therefore, in contrast to fathers, stickleback larvae respond more strongly to cues of dragonfly predators compared to sculpin, especially in freshwater populations where dragonfly larvae are endemic. This stronger response arose regardless of paternal treatment; this suggests that, consistent with a previous study [28], fathers do not induce predator-specific behavioral priming (i.e., offspring of dragonfly-exposed fathers do not respond more strongly to dragonfly larvae in their assay and vice versa for offspring of sculpin-exposed fathers; non-significant interaction between assay predator and paternal treatment for all populations for all traits).

**Figure 2:**
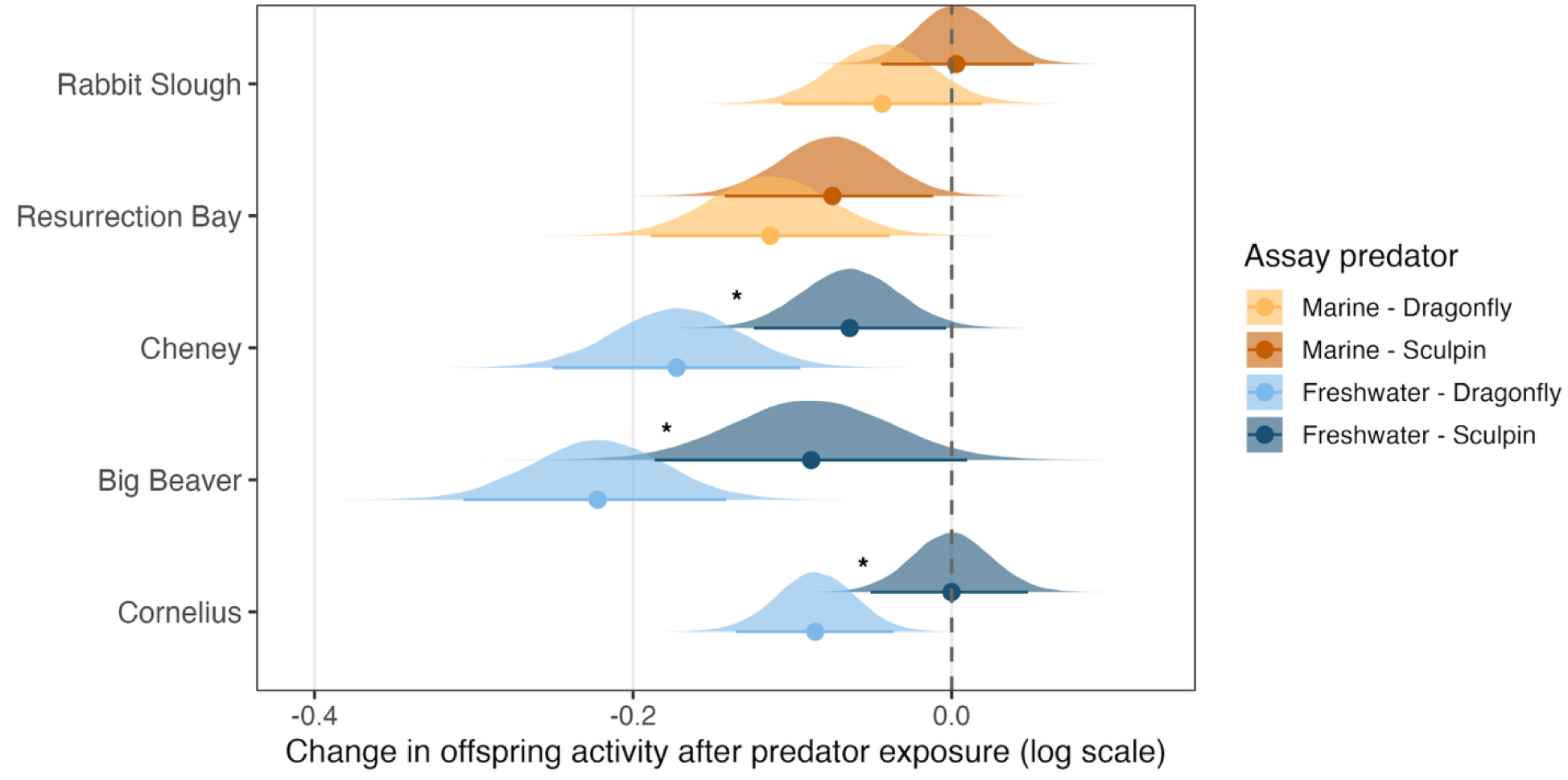
We observed offspring activity before and after a simulated predator attack by the sculpin (dark) or dragonfly (light) predator. Offspring decreased their activity after the exposure in all populations except Rabbit Slough (0 represents no change in behavior, with negative numbers indicating less activity after the simulated predator attack). Further, offspring showed a greater decrease in activity in response to dragonfly larvae compared to sculpin in freshwater populations (blue, starred), but not in marine populations (orange). Data shown are the posterior distributions of the MCMCglmm models, with the median (dot) and the 95% Highest Density Interval (bar). Black stars indicate significant differences in offspring change in activity between sculpin and dragonfly predators.

**Table 1:** Offspring traits (n = 950 across 5 populations) in response to paternal exposure to dragonfly or sculpin predators (relative to the control) for all 5 populations. Populations in orange are marine; populations in blue are freshwater. We included statistically significant interactions in the models; dashed lines indicate non-significant interactions that were not included in the final model.

|  | Activity |  |  |  |  |  |  |  |  |  |
| --- | --- | --- | --- | --- | --- | --- | --- | --- | --- | --- |
|  | Rabbit Slough (marine) |  | Resurrection Bay (marine) |  | Cheney (new FW) |  | Big Beaver (est FW) |  | Cornelius (est FW) |  |
|  | 95% CI | P | 95% CI | P | 95% CI | P | 95% CI | P | 95% CI | P |
| Dragonfly paternal treatment | 0.02, 0.35 | **0.03** | -0.46, 0.01 | 0.07 | -0.14, 0.27 | 0.51 | -0.44, 0.08 | 0.17 | -0.36, 0.17 | 0.51 |
| Sculpin paternal treatment | 0.23, 0.59 | **<0.001** | -0.71, -0.07 | **0.02** | -0.20, 0.17 | 0.94 | -0.14, 0.37 | 0.40 | -0.52, -0.05 | **0.03** |
| Offspring sex | 0.09, 0.30 | **<0.001** | -0.37, 0.03 | 0.11 | -0.11, 0.13 | 0.83 | -0.33, 0.16 | 0.50 | -0.07, 0.17 | 0.42 |
| Standard length | -0.02, 0.04 | 0.51 | 0.003, 0.09 | **0.03** | -0.02, 0.07 | 0.35 | -0.03, 0.04 | 0.64 | -0.005, 0.07 | 0.09 |
| Assay predator | -0.11, 0.11 | 0.99 | -0.02, 0.23 | 0.10 | -0.21, 0.05 | 0.23 | -0.17, 0.13 | 0.77 | -0.16, 0.07 | 0.42 |
| Time period | -0.10, 0.01 | 0.15 | -0.19, -0.04 | **0.002** | -0.38, -0.19 | **<0.001** | -0.31, -0.14 | **<0.001** | -0.14, -0.04 | **<0.001** |
| Assay predator * time period | -0.03, 0.12 | 0.25 | -0.06, 0.13 | 0.46 | 0.01, 0.20 | **0.03** | 0.01, 0.26 | **0.03** | 0.02, 0.16 | **0.02** |
| Dragonfly pat treat * sex | --- | --- | -0.08, 0.49 | 0.17 | --- | --- | 0.07, 0.76 | **0.02** | --- | --- |
| Sculpin pat treat * sex | --- | --- | 0.05, 0.63 | **0.02** | --- | --- | -0.41, 0.26 | 0.67 | --- | --- |
| Dragonfly pat treat * time | --- | --- | --- | --- | 0.09, 0.32 | **<0.001** | --- | --- | --- | --- |
| Sculpin pat treat * time | --- | --- | --- | --- | 0.03, 0.26 | **0.02** | --- | --- | --- | --- |
| Thigmotaxis |  |  |  |  |  |  |  |  |  |  |
| Dragonfly paternal treatment | -0.50, 0.13 | 0.27 | 0.28, 0.92 | **<0.001** | -0.61, 0.09 | 0.15 | -0.12, 0.62 | 0.18 | -0.59, 0.40 | 0.72 |
| Sculpin paternal treatment | -0.53, 0.07 | 0.11 | 0.04, 0.87 | **0.03** | -0.90, -0.29 | **<0.001** | -0.46, 0.22 | 0.48 | -0.40, 0.67 | 0.66 |
| Offspring sex | -0.20, 0.22 | 0.94 | -0.28, 0.11 | 0.38 | -0.03, 0.40 | 0.09 | -0.19, 0.33 | 0.64 | -0.71, 0.07 | 0.12 |
| Standard length | 0.01, 0.13 | **0.03** | -0.03, 0.11 | 0.25 | 0.03, 0.20 | **0.007** | -0.06, 0.08 | 0.76 | -0.01, 0.16 | 0.08 |
| Assay predator | -0.32, 0.12 | 0.38 | -0.33, 0.10 | 0.26 | -0.22, 0.25 | 0.91 | -0.23, 0.33 | 0.69 | -0.47, 0.05 | 0.12 |
| Time period | -0.38, -0.12 | **<0.001** | -0.27, 0 | **0.05** | -0.53, -0.22 | **<0.001** | -0.48, -0.12 | **<0.001** | -0.47, -0.10 | **0.003** |
| Assay predator * time period | -0.09, 0.28 | 0.30 | -0.25, 0.13 | 0.58 | -0.06, 0.37 | 0.14 | -0.14, 0.37 | 0.37 | -0.11, 0.27 | 0.41 |
| Dragonfly pat treat * sex | --- | --- | --- | --- | --- | --- | --- | --- | 0.20, 1.39 | **0.01** |
| Sculpin pat treat * sex | --- | --- | --- | --- | --- | --- | --- | --- | -0.59, 0.64 | 0.92 |
| Dragonfly pat treat * time | --- | --- | --- | --- | --- | --- | --- | --- | -0.21, 0.25 | 0.88 |
| Sculpin pat treat * time | --- | --- | --- | --- | --- | --- | --- | --- | -0.01, 0.45 | **0.05** |
| Freezing behavior |  |  |  |  |  |  |  |  |  |  |
| Dragonfly paternal treatment | -0.97, -0.03 | **0.04** | -0.60, 0.38 | 0.69 | -0.38, 1.21 | 0.30 | -0.13, 0.96 | 0.14 | -1.24, 0.32 | 0.24 |
| Sculpin paternal treatment | -0.85, 0.05 | 0.07 | -0.11, 0.94 | 0.11 | -0.05, 1.36 | 0.07 | -0.46, 0.65 | 0.76 | -1.43, 0.21 | 0.15 |
| Offspring sex | -0.58, 0.05 | 0.11 | -0.13, 0.54 | 0.23 | -0.34, 0.95 | 0.33 | -0.70, 0.12 | 0.16 | -0.93, 0.16 | 0.17 |
| Standard length | -0.01, 0.16 | 0.10 | -0.16, 0.07 | 0.40 | -0.34, -0.07 | **0.004** | 0.04, 0.23 | **0.005** | -0.01, 0.21 | 0.08 |
| Assay predator | -0.46, 0.15 | 0.31 | -0.66, 0.006 | **0.05** | -0.71, 0.01 | 0.06 | -1.13, -0.35 | **<0.001** | -0.32, 0.35 | 0.95 |
| Dragonfly pat treat * sex | --- | --- | --- | --- | -1.68, 0.14 | 0.10 | --- | --- | -0.24, 1.46 | 0.16 |
| Sculpin pat treat * sex | --- | --- | --- | --- | -2.14, -0.32 | **0.008** | --- | --- | 0.19, 1.91 | **0.02** |
| Standard length |  |  |  |  |  |  |  |  |  |  |
| Dragonfly paternal treatment | -0.44, 0.53 | 0.86 | -0.61, 0.40 | 0.69 | -0.96, -0.11 | **0.01** | -0.19, 1.07 | 0.19 | -0.24, 2.00 | 0.12 |
| Sculpin paternal treatment | 0.67, 2.40 | **<0.001** | 0.27, 1.60 | **0.007** | -1.06, -0.30 | **<0.001** | -0.73, 0.68 | 0.91 | -1.80, -0.12 | **0.03** |
| Offspring sex | -0.53, 0.10 | 0.19 | -0.72, -0.12 | **0.007** | -0.51, 0.04 | 0.08 | -0.71, 0.06 | 0.10 | -0.01, 0.88 | 0.055 |
|  | 0.41, 0.55 | **<0.001** | 0.56, 0.78 | **<0.001** | 0.34, 0.45 | **<0.001** | 0.52, 0.70 | **<0.001** | 0.33, 0.46 | **<0.001** |
| Dragonfly pat treat * sex | --- | --- | --- | --- | --- | --- | --- | --- | -1.64, -0.25 | **0.009** |
| Sculpin pat treat * sex | --- | --- | --- | --- | --- | --- | --- | --- | -1.35, 0.01 | 0.054 |
| Body condition |  |  |  |  |  |  |  |  |  |  |
| Dragonfly paternal treatment | 0.00001, 0.005 | **0.04** | 0.001, 0.007 | **0.002** | -0.005, 0.002 | 0.52 | -0.006, -0.001 | **0.005** | -0.01, 0.01 | 0.99 |
| Sculpin paternal treatment | -0.006, 0.007 | 0.85 | -0.002, 0.005 | 0.30 | -0.002, 0.003 | 0.68 | -0.007, -0.002 | **<0.001** | -0.008, 0.002 | 0.27 |
| Offspring sex | -0.002, 0.0009 | 0.42 | -0.002, 0.001 | 0.81 | -0.001, 0.002 | 0.70 | -0.002, 0.001 | 0.62 | -0.002, 0.002 | 0.99 |
| Season (date measured) | -0.0001, -0.00005 | **<0.001** | -0.00004, 0.0001 | 0.40 | -0.00004, 0.0004 | 0.27 | -0.0002, -0.00006 | **<0.001** | -0.00002, 0.0004 | 0.08 |

### No evidence for variation in the strength of TGP between ancestral marine and derived freshwater populations

Given evidence for parallel evolution of plastic antipredator responses in freshwater populations, we then tested for three signatures of adaptive evolution of TGP due to paternal predation exposure. First, we tested the flexible stem hypothesis by assessing if TGP is genetically assimilated in derived freshwater populations. We found limited evidence for variation in the strength of TGP between ancestral marine and derived freshwater populations (n = 950 offspring tested across 5 populations): paternal exposure to sculpin induced main or interactive effects in 5/10 traits across the two marine populations, in 4/5 traits in the one newly derived freshwater population, and in 5/10 traits across the two established freshwater populations. Similarly, paternal exposure to dragonfly larvae induced main or interactive effects in 5/10 traits across the two marine populations, in 2/5 traits in the one newly derived freshwater population, and in 4/10 traits across the two established freshwater populations. Collectively, these results suggest that freshwater populations retain the ability to mount strong transgenerational responses to both predators and that marine populations show strong TGP even in response to novel dragonfly predators.

### Evidence for parallel effects of paternal predation exposure on offspring growth, but not on behavior

Because we found TGP in all populations and for both predator types, we then tested to see if this variation had adaptive signatures by testing for 1) parallel evolution of TGP across marine and freshwater populations and 2) fine-tuning of TGP, resulting in more adaptive or less variable TGP in response to endemic predators compared to novel predators. In both marine populations, paternal exposure to predation risk enhanced growth: offspring of dragonfly-exposed fathers were in better body condition (higher mass relative to length) compared to offspring of control fathers, while offspring of sculpin-exposed fathers were longer in both marine populations (Table 1; Figure 3). In contrast, paternal exposure to predation risk largely slowed growth in freshwater populations. In Big Beaver, offspring of both sculpin and dragonfly-exposed fathers were in worse body condition relative to offspring of control fathers while in Cheney, offspring of both sculpin and dragonfly-exposed fathers were shorter relative to offspring of control fathers (Table 1; Figure 3). In Cornelius, offspring of sculpin-exposed fathers were shorter relative to offspring of control fathers; this was also true for sons of dragonfly-exposed fathers, although daughters of dragonfly-exposed fathers were longer (main and sex-specific effects of paternal treatment; Table 1). Because faster growth is usually adaptive by allowing individuals to access more food resources and outgrow the gape of the predator earlier in development [36], these results suggest that TGP in response to predators seems more likely to be adaptive in marine populations where dragonfly are novel compared to freshwater populations where dragonfly are endemic.

**Figure 3:**
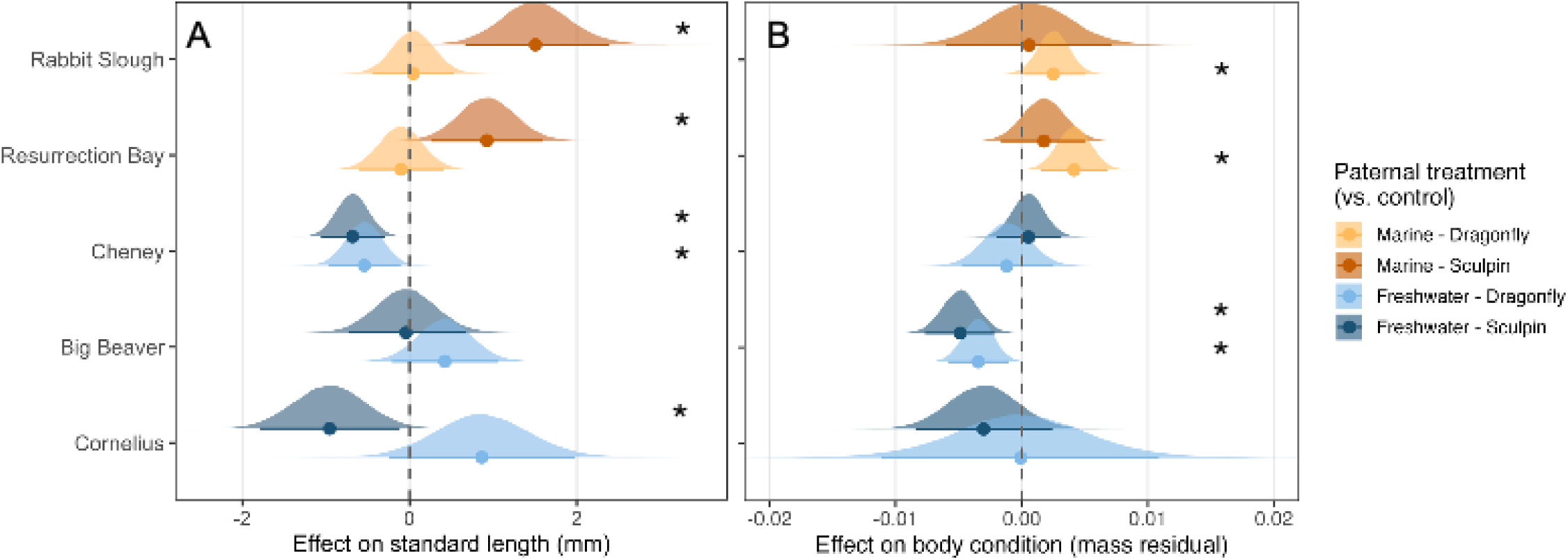
Length (A) and body condition (mass, controlling for length; B) in marine (orange) and freshwater (blue) populations in response to paternal exposure to dragonfly larvae (light), or sculpin predator (dark) treatment. Data shown are the posterior distributions of the MCMCglmm models, with the median (dot) and the 95% Highest Density Interval (bar); bars that do not overlap zero indicate that the paternal treatment effect is significantly different than the control. Black stars indicate significant main effects of paternal treatment; note that, for length, there is a significant sex-specific of paternal exposure to dragonfly larvae in Cornelius that is not visualized here (sons of dragonfly-exposed fathers were shorter while daughters were longer).

In contrast to growth, we found no evidence for parallel paternal effects on offspring behavior. For example, changes in offspring behavior induced by paternal treatment were largely opposite between the two marine populations. In Rabbit Slough, offspring with predator-exposed fathers were bolder (more active and froze less relative to offspring of control fathers; Table 1; Figure 4). In contrast, in Resurrection Bay, offspring with predator-exposed fathers were less bold (less active and higher thigmotaxis relative to offspring of control fathers, with the decrease in activity for sculpin-exposed fathers stronger in daughters than sons (main and interactive effects of paternal treatment; Table 1; Figure 4). Similarly, the effect of paternal treatment on offspring behavior was highly variable across the three freshwater populations (Figure 4; Supplementary Figure 1; see below), suggesting that offspring traits reflect a combination of parallel evolution and stochasticity.

**Figure 4:**
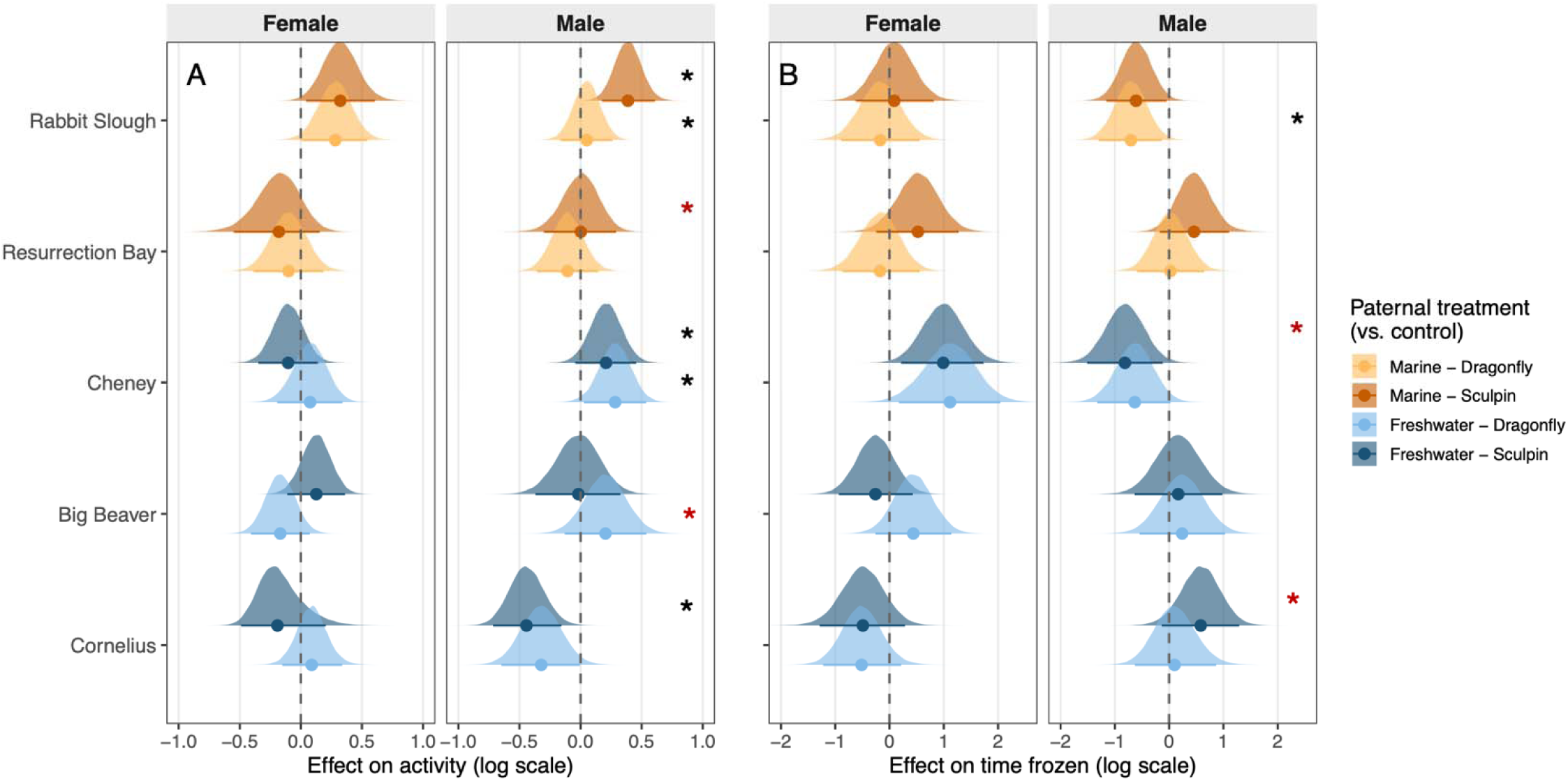
Variation in offspring activity (A) and freezing time after a simulated predator attack (B) in marine (orange) and freshwater (blue) populations due to paternal exposure to dragonfly larvae (light), or sculpin predator (dark) treatment. Data shown are the posterior distributions of the MCMCglmm models, with the median (dot) and the 95% Highest Density Interval (bar); bars that do not overlap zero indicate that the paternal treatment effect is significantly different than the control. Black stars indicate significant main effects of paternal treatment, while red stars indicate significant interactions between paternal treatment and offspring sex. Note that, for activity, there is a significant paternal treatment by time interaction in Cheney that is not visualized here (offspring of dragonfly and sculpin-exposed fathers showed a reduced change in activity in response to the simulated predator attack compared to offspring of control fathers).

In addition to mean shifts in offspring traits, examining variance in offspring traits can reveal if TGP refines reaction norms in response to endemic predators. If there is high interindividual variation in response to novel dragonfly in marine populations, we would expect that paternal exposure to dragonfly larvae would increase variance in offspring traits in marine populations, but not in freshwater populations. We found limited evidence for this possibility: paternal treatment to dragonfly larvae did not increase trait variance in one marine population (Rabbit Slough). In the other marine population (Resurrection Bay), paternal exposure to both dragonfly larvae and sculpin increased variance of offspring thigmotaxis and length relative to the control (Supplementary Table 4), suggesting that this shift in variance occurs in response to both endemic and novel predators.

### Sex-specific effects in offspring are linked to paternal exposure to endemic predators

Although we did not find fine-tuning consistent with our initial predictions, we did find evidence that sex-specific effects are linked to endemic predators. We found no sex-specific responses of paternal exposure to novel dragonfly larvae in marine populations or the newly derived freshwater population, although we did find sex-specific responses of paternal exposure to endemic sculpin in Resurrection Bay (activity; Table 1; Figure 4A). However, we did find strong sex-specific responses of paternal exposure to both dragonfly larvae and sculpin in established freshwater populations. In Big Beaver, offspring of dragonfly-exposed fathers showed sex-specific changes in activity (sons were more active and daughters were less active relative to the controls; Table 1; Figure 4A). In Cornelius, offspring of dragonfly-exposed fathers showed sex-specific changes in thigmotaxis (sons, but not daughters, were more anxious relative to the controls) and length (sons of dragonfly-exposed fathers were shorter while daughters were longer (Table 1)). Further, in Cornelius, offspring of sculpin-exposed fathers showed sex-specific changes in freezing behavior (sons, but not daughters, spent more time frozen relative to controls; Table 1; Figure 4B). Collectively, these results suggest that, despite high variability in the effects of paternal treatment on offspring behavior, these effects are often sex-specific in response to endemic predators.

## Discussion

Because plastic responses to new environments can alter genetic adaptation by moving traits closer to or farther from their new local optimum (Levis & Pfennig, 2016), understanding the evolution of TGP can provide critical insight into the mechanisms allowing organisms to persist in novel environments. Here, we measured sperm-mediated paternal effects across ancestral marine and derived freshwater stickleback populations to test if TGP shows signatures of an adaptive evolutionary process, with systematic variation among populations that is related to population type or a population’s evolutionary history with a specific predator. We had three nonexclusive adaptive hypotheses. First, the flexible stem model of adaptive evolution predicts consistent differences in the extent of plasticity between ancestral versus derived populations. We found no evidence for this: there were no systematic differences in the magnitude of TGP between marine and freshwater populations in response to either predator. Second, if TGP is adaptive, we expect to see signatures of parallel evolution. We found some evidence consistent with parallelism in both within and trans-generational plasticity: 1) personal exposure of larval stickleback to dragonfly larvae elicited conserved antipredator responses in freshwater populations that were absent in marine populations, and 2) paternal predation exposure enhanced offspring growth in marine populations while slowing growth in freshwater populations. Third, if TGP is adaptive, then we expect to see evidence that TGP is fine-tuned. We found some evidence for fine-tuning with respect to sex-specific TGP: endemic predators elicited sex-specific responses in offspring (especially in freshwater populations), while non-endemic predators did not. Collectively, our results agree with previous studies documenting extensive interpopulation variation in TGP (Walsh et al., 2016), with this variation reflecting a combination of adaptive and stochastic processes.

Paternal exposure to both dragonfly and sculpin predators altered offspring traits in all populations, providing limited evidence that plasticity in canalized in derived freshwater populations. Instead, we found strong, but variable TGP, across populations. Some of this variation was systematic. There was evidence for parallel evolution with respect to body size: paternal exposure to predators enhanced offspring growth in marine populations and slowed growth in freshwater populations. Interestingly, these effects were present even in Cheney, which was experimentally seeded from Rabbit Slough only 8 years before sampling, suggesting these effects can evolve rapidly. In contrast, shifts in behavioral traits seemed less consistent: the type of offspring behavior that was altered and direction of the change in behavior was sometimes completely opposite within the same population type. This is consistent with previous studies in fish that have found mixed or weak evidence for parallel evolution among replicate populations (Bolnick et al., 2018). Environmental heterogeneity among replicate marine and freshwater population can contribute to nonparallel evolution, especially in behavior where factors such as habitat complexity, predator density, or other predation threats (e.g., birds) can have a strong influence on the optimal behavioral response to predation (e.g., hide if dense vegetation, flee if open) (Hendry et al., 2024). Further, a portion of variation in TGP may have been induced due to grandparental effects and differences in the starting genotypes among populations, potentially limiting our ability to fully detect parallel evolution.

We also tested for evidence for the fine-tuning hypothesis, or how the evolutionary history of a population with a stressor influences the strength and adaptive nature of TGP. Contrary to the fine-tuning hypothesis, we found little evidence for more adaptive or less variable TGP in freshwater populations where dragonfly are endemic compared to marine populations where dragonfly are novel. Instead, there were potentially adaptive increases in offspring size when fathers were exposed to dragonfly larvae in marine populations, with putatively less adaptive growth effects in freshwater populations where dragonfly larvae are endemic. These results are in contrast to other studies that have found negative carryover effects of parental experiences in populations where parental cues are novel compared to populations where cues are endemic (Neylan et al., 2022; Sultan et al., 2009). However, given that predation risk is a salient stressor in all stickleback populations, individuals in marine populations may already be primed to respond to a variety of stressors they perceive as predator threats. Providing support for this, although paternal exposure to dragonfly larvae often induced different traits than paternal exposure to sculpin, we also found that when both paternal exposure to sculpin and dragonfly elicited effects on the same offspring trait, they were always in the same direction within a population (e.g., offspring with dragonfly or sculpin-exposed fathers were more anxious (less active with higher thigmotaxis) relative to offspring of control fathers in Resurrection Bay). Therefore, high predation risk in all our populations (e.g., from birds, piscivorous fish) may have selected for TGP that induces a more general phenotype that broadly prepares offspring for stressful environments (Chen et al., 2021; Potticary & Duckworth, 2020; Sandner et al., 2018). This may be adaptive if parents cannot accurately identify the stressor they encounter or if a common mechanism (e.g., glucocorticoids) underlies the response to environmental stress (Donelan et al., 2020).

Although we found limited evidence for our initial fine-tuning predictions, we did find strong sex-specific effects of paternal exposure to endemic predators. Although more work is needed to see if these results hold across a greater number of populations, these results open up the possibility that sex-specific transgenerational plasticity may be evolving across populations, with these sex-specific responses organizing early in offspring development well before sex-specific life history strategies emerge. Previous work that has found that parental environments can have opposing effects on the same trait in sons compared to daughters or can influence different traits in sons versus daughters (A. M. Bell & Hellmann, 2019), with little evidence that offspring attend primarily to cues from their same-sex parent (i.e., stronger paternal effects in sons compared to daughters) (Hellmann, Bukhari, et al., 2020). It is possible that tailored responses could enhance the adaptive nature of TGP by allowing offspring to differentially respond to paternal cues depending on sex-specific life history strategies and/or vulnerability to ecological stressors; however, more tests are needed to determine if sex-specific TGP represents adaptive evolution or simply mechanistic differences in how sons and daughters process paternal cues during early embryonic development (e.g., sex-biased gene expression) (Hellmann, Bukhari, et al., 2020). Nevertheless, if we had neglected to measure offspring sex in these larval offspring, we would have failed to detect many of our significant findings, suggesting that TGP may be underreported in studies that fail to measure and test for the influence of offspring sex.

Our study exposed fathers to two different predator types: dragonfly larvae and sculpin fish. Interestingly, although fathers did not behaviorally respond to the dragonfly larvae, we found strong transgenerational effects of paternal exposure to dragonfly larvae. Given that dragonfly larvae are juvenile predators, this suggests that stimuli may not need to be threatening to fathers to elicit an effect in offspring. This is consistent with TGP representing adaptive information priming: parents reliably assess their environment (e.g., level of predation risk, competition) and provide offspring with relevant information about their future environment. Given that juveniles themselves responded more strongly to dragonfly larvae, especially in freshwater populations where they are endemic, fathers may be transmitting important information about pressing threats offspring might encounter immediately after hatching. However, future studies investigating the fitness consequences of these trait changes are needed to assess whether these shifts in offspring phenotypes are adaptive.

## Conclusions

We designed a robust experiment-rearing individuals in a common garden and investigating TGP across multiple populations and predators- to thoroughly test for evidence of adaptive evolution of TGP. Our results demonstrate that there are strong sperm-mediated paternal effects across multiple populations of sticklebacks, suggesting that paternal experiences substantially alter offspring phenotypes even in the absence of physical interactions between fathers and offspring (i.e., paternal care). We found a high degree of plasticity across all population, with signatures of both parallel and nonparallel evolution. Further, there was evidence that sex-specific offspring traits emerged primarily in relation to endemic predators, indicating that evolutionary history may shape more targeted plastic responses. These results suggest that ecological differences among populations may shape TGP in a way that reflects an adaptive evolutionary process, but that there is also substantial variation that may reflect stochasticity or environmental heterogeneity that we did not account for. However, adaptive evolution is a two-step process, in which heritable genotypic and phenotypic variation must first be present and then selected on (West-Eberhard, 2003). While this stochasticity may be random or non-adaptive in many circumstances, this standing phenotypic variation is also necessary for evolution (Levis & Pfennig, 2016). More research, integrating tightly controlled lab experiments with field experiments in a natural setting, could determine the extent to which the ecological context influences how environmental cues are integrated into parental phenotypes and whether standing variation in TGP leads to adaptive consequences for offspring traits.

